# Dendritic GABA_B_ receptors control nonlinear information transfer along the dendro-somatic axis in layer 5 pyramidal neurons

**DOI:** 10.1101/762625

**Authors:** Jan M. Schulz, Matthew E. Larkum

**Author notes:** **Correspondence:** Dr Jan M. Schulz, Department of Biomedicine, University of Basel, Pestalozzistr. 20, 4056 Basel, Switzerland.

## Abstract

Dendritic GABA_B_ receptors (GABA_B_Rs) mediate a slow form of interhemispheric inhibition. Surprisingly, this inhibition has no detectable effect on the somatic membrane potential of layer 5 pyramidal neurons, whereas the action potential (AP) output is robustly decreased even when the input is proximal to the cell body. To elucidate the underlying mechanisms, we systematically mapped the AP frequency-current (F-I) relationship during dual patch-clamp recordings from soma and apical dendrite. The AP output function was governed by the synergistic interaction between dendritic and somatic compartments as the local input and transfer resistance from dendrite to soma (K_ds_) depended on the dendritic membrane potential. Thus, K_ds_ doubled at an estimated rate of once per 28.7 mV depolarization due to HCN channel deactivation. In addition, dendritic L-type Ca^2+^ channels converted individual APs into dendritic Ca^2+^ spikes causing high-frequency bursts of APs (HFB) during large dendritic depolarization. Activation of dendritic GABA_B_Rs greatly reduced both nonlinear mechanisms. While direct block of L-type Ca^2+^ channels reduced the number of HFBs, K^+^ channel activation decreased voltage-dependent input and transfer resistances and decreased the AP rate under all conditions. These results highlight the powerful modulation of the input integration in pyramidal neurons by metabotropic receptor-activated K^+^ channels.

## INTRODUCTION

The mammalian cortex is a conspicuously layered structure that receives inputs from afferent subcortical and other cortical areas in a layer-specific manner (Douglas & Martin, 2004). The dendrites of pyramidal neurons typically span several layers and can therefore integrate inputs from different afferent sources. Together with the neurons’ intrinsic physiological properties, this anatomical arrangement places pyramidal neurons in the ideal position to form associative computational elements of different input streams (Larkum, 2013). It has been suggested that pyramidal neurons, in particular those residing in layer 5 (L5), act as coincidence detectors of sensory feedforward inputs impinging onto the basal dendrites and higher-order feedback (e.g. attentional) inputs onto the apical tuft (Xu *et al.*, 2012; Takahashi *et al.*, 2016). One important mechanism for coincidence detection is the generation of dendritic Ca^2+^ spikes by the interaction of backpropagating APs and large dendritic postsynaptic potentials (PSP) resulting in a burst of axonal APs (Larkum *et al.*, 1999). This mechanism also contributes to the amplification of AP output during noisy somatic input currents by additional dendritic depolarization, i.e. it increases the gain of the input-AP output (F-I) relationship (Larkum *et al.*, 2004). Although other cell-intrinsic voltage-dependent mechanisms, such as the hyperpolarization and cyclic nucleotide activated (HCN) cation current (I_H_), are known to shape synaptic integration (Williams & Stuart, 2000; Berger *et al.*, 2001; Harnett *et al.*, 2015), their contributions to the synergistic interaction between depolarization in the apical dendritic and perisomatic compartments are less well understood (Phillips *et al.*, 2016).

Active dendritic processing significantly contributes to sensory perception and behaviorally relevant neuronal computations (Xu *et al.*, 2012; Harnett *et al.*, 2013; Smith *et al.*, 2013; Bittner *et al.*, 2015; Takahashi *et al.*, 2016; Sheffield *et al.*, 2017). Not surprisingly, the integrative properties of pyramidal neurons are the target of multiple modulatory systems including serotonergic (Santello & Nevian, 2015), adrenergic (Liu *et al.*, 2017) and cholinergic signals (Williams & Fletcher, 2018). Thus, cholinergic inputs from the basal forebrain involved in attentional processes have recently been shown to enhance dendritic R-type Ca^2+^ channel activity via muscarinic receptor activation in L5 pyramidal neurons (Williams & Fletcher, 2018). In contrast, down-modulation of dendritic excitability by GABA_B_ receptors (GABA_B_Rs) involves both direct inhibition of dendritic L-type Ca^2+^ channels and activation of G protein-coupled inwardly-rectifying K^+^ (GIRK) channels (Perez-Garci *et al.*, 2006; Palmer *et al.*, 2012; Perez-Garci *et al.*, 2013). GABA_B_Rs mediate a slow form interhemispheric inhibition that decreases the *in vivo* AP rate specifically in L5 pyramidal neurons by about a third (Palmer *et al.*, 2012; Palmer *et al.*, 2013). This form of interhemispheric inhibition acts as silent inhibition that reduces L5 pyramidal neuron spiking without noticeable effects on the subthreshold voltage envelope at the cell body. While there is strong evidence that direct L-type Ca^2+^ channel inhibition fundamentally changes the integrative properties of pyramidal neurons (Perez-Garci *et al.*, 2006; Breton & Stuart, 2012; Palmer *et al.*, 2012; Perez-Garci *et al.*, 2013), the relative contribution of dendritic K^+^ channels to the decreased somatic AP output has remained unclear.

In the current study, we sought to elucidate the precise contributions by both GABA_B_R-activated effector pathways to the modulation of integrative properties of L5 pyramidal neurons.

## RESULTS

We systematically mapped the F-I relationship and membrane depolarization during combined current injections into the somatic and apical dendritic compartments of thick-tufted L5b pyramidal neurons in rat (P28-40) somatosensory cortex using dual patch-clamp recordings. Somatic current injection into L5b pyramidal neurons resulted in classic type 1 action potential (AP) firing (Hodgkin, 1948). The frequency-input (F-I)-curve was well fitted by a square root function (see equation (*1*) in Methods; Fig. 1A-B). To include the effect of dendritic current injections, we added the dendritic current scaled by a factor *D* in a passive dendrite spike rate model (PDSR model described by equation (*2*) in Methods; Fig. 1C). If the dendrite behaves like a passive cable, any dendritic current depolarizes the soma according to the transfer resistance from dendrite to soma (K_ds_= V_soma_/I_dend_). Hence, the scaling factor *D* was set to the ratio of transfer resistance relative to the somatic input resistance measured at resting membrane potential (*D*=K_ds_/R_in_). However, the experimentally observed F-I relationship during combined dendritic and somatic current injections exhibited considerably higher AP rates than predicted by the PDSR model (Fig.1E and F). To determine the parameters that caused this deviation, we fitted the model (equation (*2*) to the data (Fig. 1J) and compared the obtained parameters to those of the original PDSR model. The obtained gain and current threshold (I_T_) were very similar between fits to purely somatic compared to combined somatic and dendritic current injections (Fig. 1L). However, the experimentally established *D* (0.48 ± 0.03, n=25 neurons) was significantly larger than K_ds_/R_in_ (0.29 ± 0.02, *P*<0.0001; Wilcoxon signed rank test). The strong nonlinear effects of dendritic current injections on spike patterns that included dendritic Ca^2+^ spikes and somatic burst firing (Fig. 1D) clearly indicated that active dendritic mechanisms significantly contributed to the F-I relationship.

**Figure 1.**
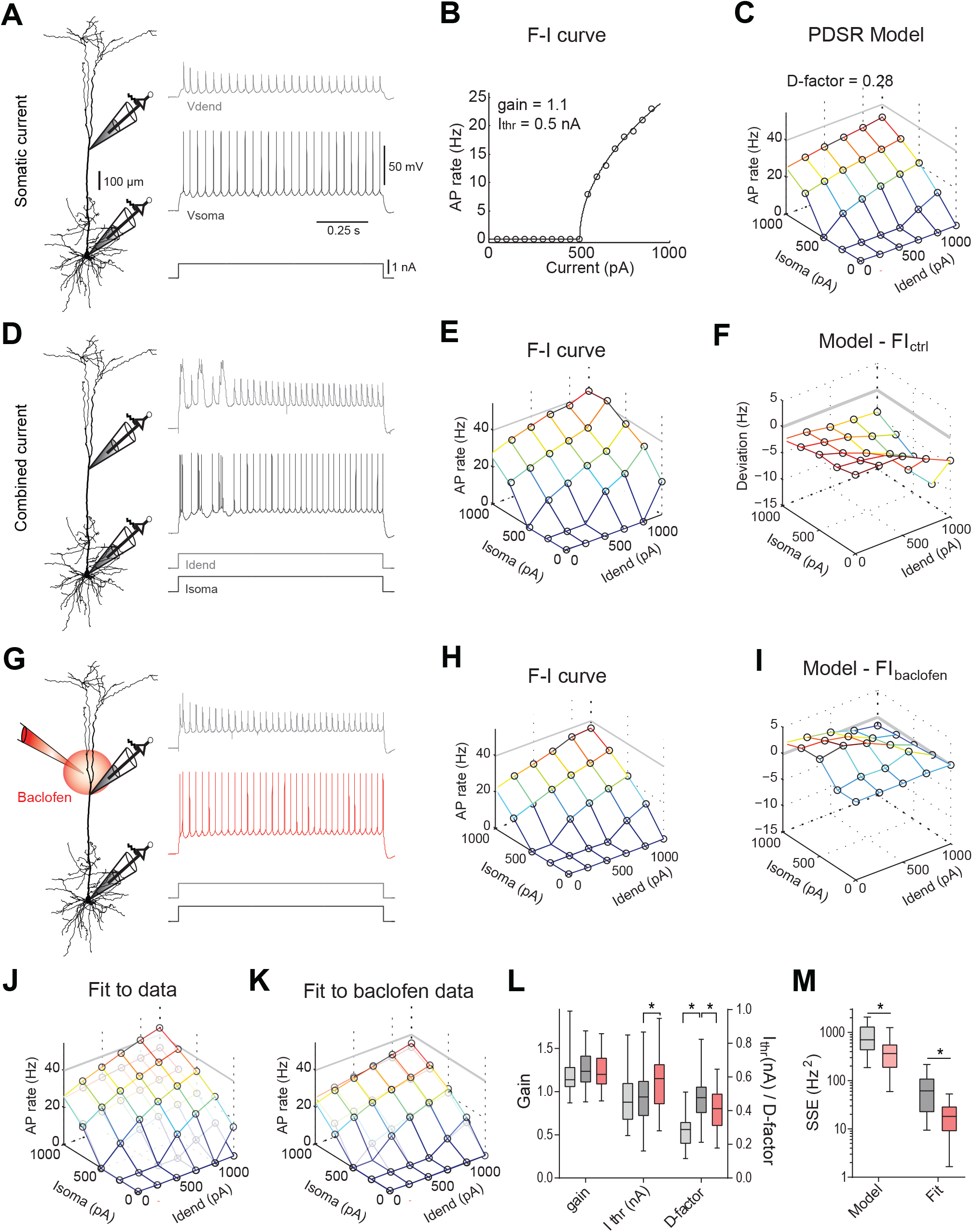
The contribution of active dendritic processes to the F-I relationship is minimized by dendritic GABA_B_R activation. **A**) Left, locations of dual dendritic and somatic patch-clamp recordings are indicated on a biocytin-filled layer 5 pyramidal neuron. Dendritic and somatic membrane potential responses to a current step in the somatic patch pipette. **B**) AP rate versus current (F-I) relationship for somatic current injections. Parameters of eq. (*1*) from the fit to the data of the same neuron are indicated. **C**) Extrapolation of the F-I relationship for combinations of dendritic and somatic current injections according to the passive dendrite spike rate (PDSR) model in eq. (*2*) with *D*=K_ds_/R_in_. **D**) Membrane potential responses of the same neuron to combined dendritic and somatic current injection. **E**) Experimentally observed F-I relationship. **F**) The average difference of the experimental F-I relationship from the neuron-specific PDSR model (n=23 neurons) shows a strong deviation from the passive case indicating a pronounced contribution of active dendritic processes to the F-I relationship. **G**) Membrane potential responses to combined dendritic and somatic current injection during a puff of baclofen onto the apical dendrite. **H**) Experimentally observed F-I relationship for the same neuron. **I**) During the baclofen puff, the average deviation of the F-I relationships from the PDSR model is small (n=23 neurons). **J**) Fit of eq. (*2*) to the experimental data obtained from the same neuron during combined dendritic and somatic current injection (see Fig. 1E). Note the larger AP rates compared to the PDSR model shown in lighter colors for comparison. **K**) Fit to the F-I data recorded from the same neuron during a puff of baclofen onto the apical dendrite in H. Note the striking similarity to the PDSR model shown in lighter colors. **L**) Group data (n=25) of parameters of the PDSR model (light gray) and the fits of control data (gray) and baclofen data (red) to eq. (*2*). Statistically significant differences are indicated (*, P<0.01, Wilcoxon signed rank test). **M**) Summed squared errors (SSE) of experimental F-I data to the PDSR model (left) and to the fit of eq. (*2*) (right). Data during baclofen puff (red) showed significantly lower SSE (both: *P*<0.01, Wilcoxon signed rank test, n=23). Abbr.: Idend, dendritic current; Isoma, somatic current; Vdend, dendritic membrane potential; Vsoma, somatic membrane potential

### Dendritic GABA_B_Rs minimize active dendritic contribution to the F-I relationship

Modulation via dendritic GABA_B_Rs is known to suppress dendritic Ca^2+^ spikes in L5 pyramidal neuron (Perez-Garci *et al.*, 2006; Palmer *et al.*, 2012; Perez-Garci *et al.*, 2013). We tested the effect of dendritic GABA_B_R activation on the F-I relationship by puffing baclofen (50 μM) directly onto the apical dendrite (Fig. 1G). As previously documented (Perez-Garci *et al.*, 2006; Palmer *et al.*, 2012), baclofen prevented the activation of dendritic Ca^2+^ spikes and decreased the overall AP output (Fig. 1H). The F-I relationship during dendritic GABA_B_R activation was surprisingly similar to the PDSR model’s predictions (Fig. 1I and K). Thus, the summed squared error (SSE) of the experimentally observed F-I relationship to the PDSR model was reduced (*P*=0.0031, Wilcoxon signed rank test, n=23; Fig. 1M left). Similarly, SSE of the data fit to equation (2) was reduced indicating greater qualitative similarity of the actual F-I relationship to equation (*2*) (*P*<0.0001, Wilcoxon signed rank test, n=23; Fig. 1M right). Furthermore, this fit showed a significant reduction of *D* (*P*=0.0023, Wilcoxon signed rank test, n=25) and increase of the current threshold during dendritic GABA_B_R activation (*P*<0.0001; Fig. 1L). Together, these results showed that dendritic GABA_B_Rs removed active dendritic contributions and established an F-I relationship closely resembling the PDSR model.

### Dendritic GABA_B_ modulation reduces nonlinearities at subthreshold membrane potentials

We next tested the effect of dendritic GABA_B_R activation on subthreshold membrane potential deflections in the absence of AP-associated dendritic Ca^2+^ spikes. Families of increasing current steps were injected into either dendrite or soma before and after puffing baclofen locally onto the apical dendrite (Fig.2A and B). When current steps of increasing amplitude were injected into the dendrite, the dendritic as well as the somatic membrane depolarizations increased in a supralinear fashion. Thus, the apparent local input resistance at the dendrite increased with increasing current step size (Fig. 2C, black). This effect was even stronger for the transfer resistance K_ds_ (Fig. 2D, black). Dendritic GABA_B_R activation strongly suppressed this nonlinear increase of both the local input resistance in the dendrites (*P*=0.0017; extra sum-of-squares F test, F(1,110)=10.38; Fig.2C, orange) and the transfer resistance (*P*<0.0001; F(1,236)=35.07; Fig.2D, orange). In addition, current steps of increasing amplitude into the soma induced supralinearly increasing membrane depolarizations in the soma. While dendritic GABA_B_R activation had no effect on the somatic input resistance measured at resting membrane potential (Fig. 2E), it did reduce the nonlinear increase of the somatic input resistance significantly (*P*=0.0008; extra sum-of-squares F test, F(1,119)=11.82). These effects may explain the previously made observation that dendritic GABA_B_R activation acts as silent inhibition without an apparent effect on the somatic input resistance (Palmer *et al.*, 2012). To better understand the nonlinearities in the integrative properties of L5 pyramidal neurons, we proceeded to map systematically the current-voltage relationship at subthreshold membrane potentials.

**Figure 2.**
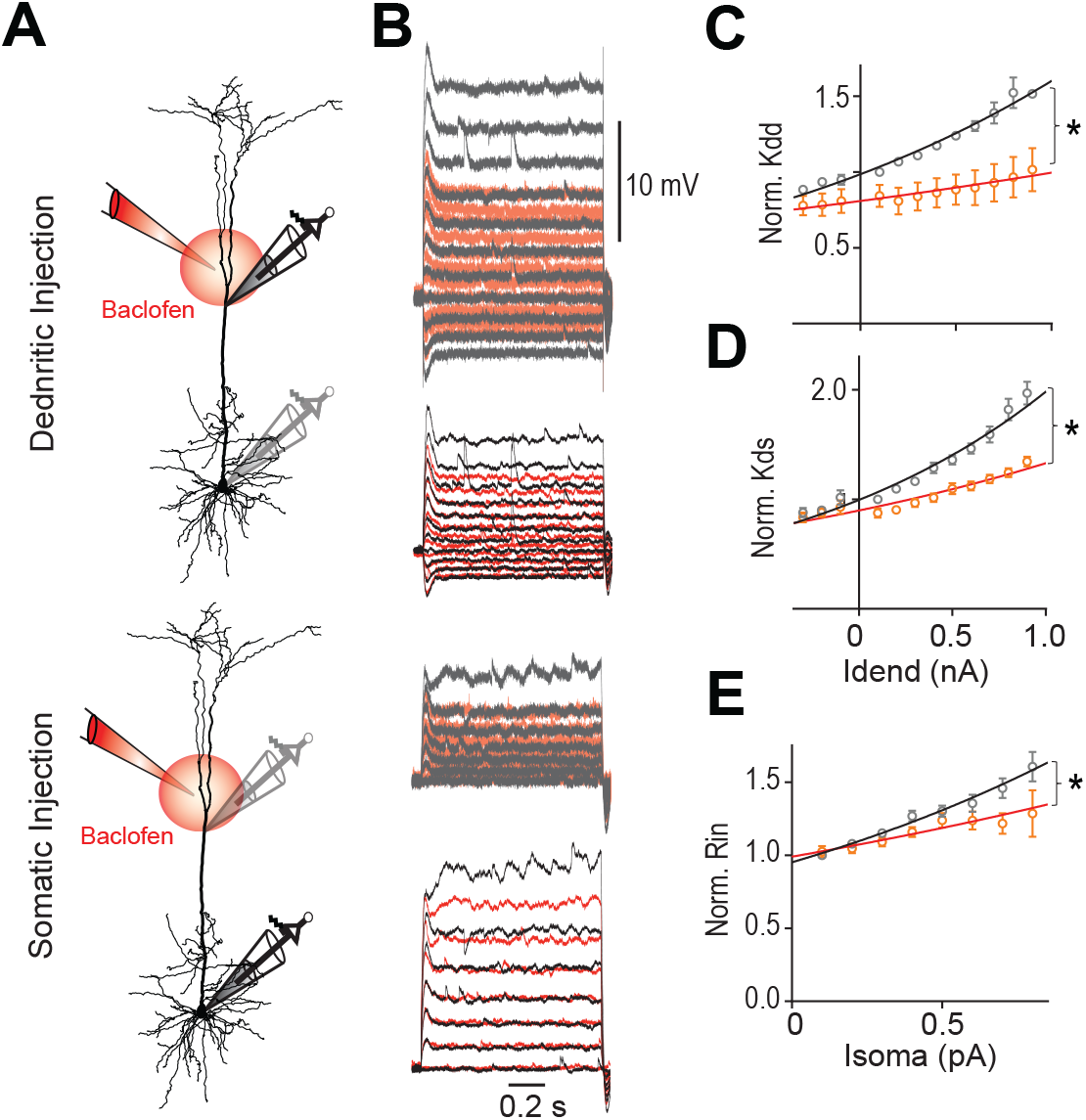
GABA_B_R activation reduces nonlinearities of input and transfer resistance. **A**) Schematic of recording arrangement during dual dendritic and somatic patch-clamp recordings. **B**) Families of current steps were injected into the dendrite (top) or soma (bottom) to evoke subthreshold membrane potential responses before (grey) and during the local application of baclofen (red) onto the apical dendrite. **C**) Local dendritic input resistance (K_dd_) normalized to values at resting membrane potential are plotted versus dendritic current amplitude. Solid line indicates exponential fit (y=*Y0*\*e(*k*\*x)) to the data. Baclofen application significantly decreased *k* (*P*=0.0017, n=5 neurons, F(1,110)=10.38), thereby increasing the doubling interval from 1.4 to 3.3 nA. **D**) Dendro-somatic transfer resistance (K_ds_) normalized to values at resting membrane potential are plotted versus dendritic current amplitude. Baclofen application significantly decreased *k* of the exponential fit indicating an increase of the doubling interval from 1.0 to 1.8 nA of K_ds_ (*P*<0.0001, n=11, F(1,236)=35.07). **E**) Somatic input resistance (R_in_) normalized to values at resting membrane potential are plotted versus somatic current amplitude. Baclofen application significantly increased the doubling interval of Rin from 1.1 to 1.9 nA (*P*=0.0008, n=11, F(1,119)=11.82).

### The transfer resistance is voltage-dependent

In linear systems, the transfer resistance is independent of the direction in which it is measured, i.e. it is transitive. In agreement with this prediction, K_ds_ closely matched K_sd_ (Fig. 3A and B). In addition, the transfer resistance is expected to depend on the distance between the two recordings sites and to decrease with increasing distance. Indeed, K_ds_ normalized to the somatic R_in_ showed a monoexponential decay with a length constant of 383 μm (Fig. 3C). However, larger current steps injected into the dendrite resulted in supralinear increases of the somatic membrane potential fluctuations (Fig. 3A). Plotting the observed K_ds_ normalized by the K_ds_ measured at the resting membrane potential versus the steady-state membrane potential locally induced by the current step showed that K_ds_ strongly depended on the membrane potential (Fig. 3D). On average, depolarization of 28.7 mV in the dendrite caused the K_ds_ to increase by a factor of two (fitted line). The local dendritic input resistance (K_dd_) was also modulated by the dendritic membrane potential (Fig. 3E); however, this voltage-dependence consisted of a doubling every 59.5 mV which was a significantly lower rate than for K_ds_ (*P*<0.001; extra sum-of-squares F test, F(3,783)=90.35).

**Figure 3.**
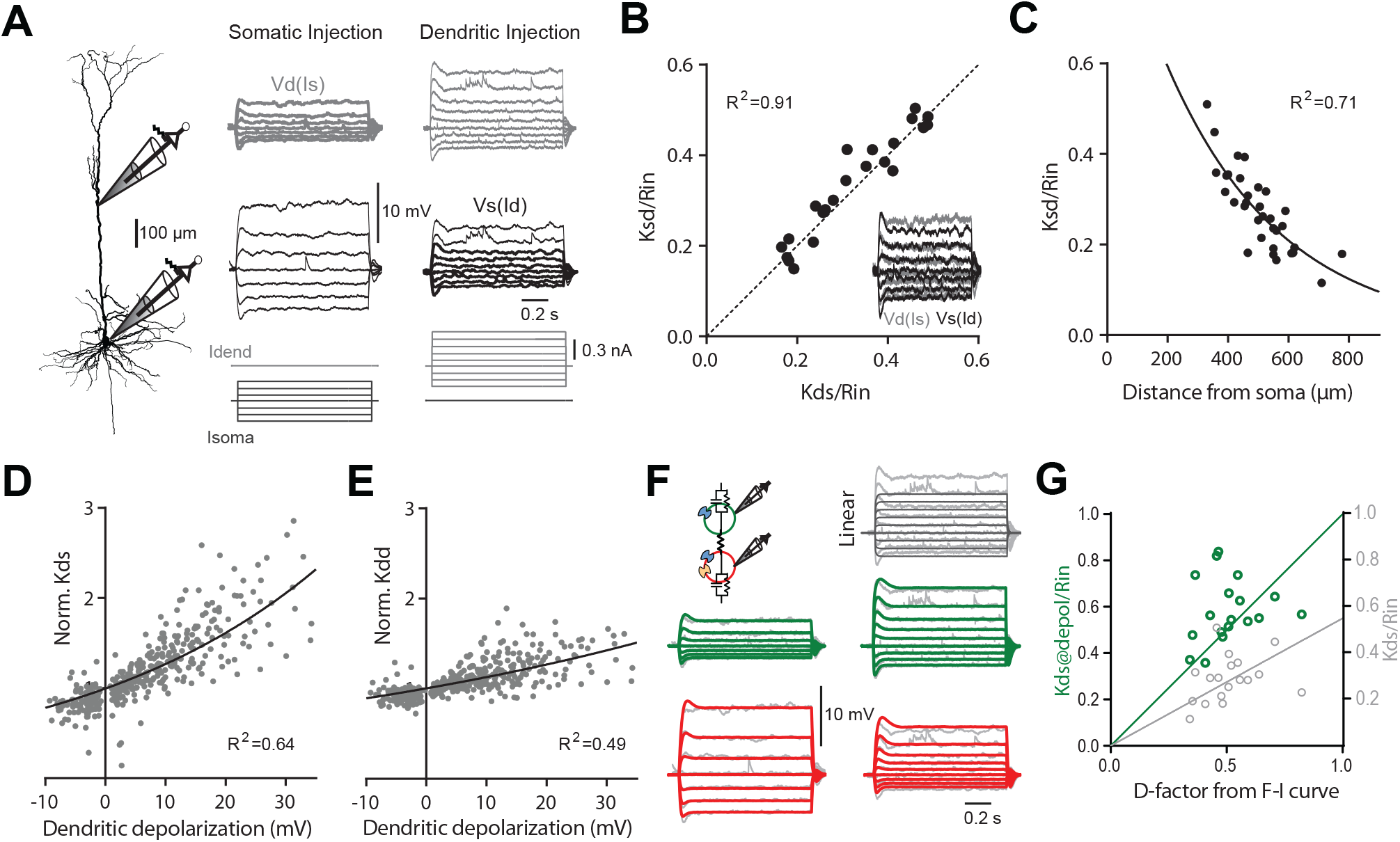
The transfer and dendritic input resistances depend on the membrane potential. **A**) Left, schematic of recording arrangement during dual dendritic and somatic patch-clamp recordings. Families of current steps were injected into the soma (center) or the dendrite (right) to evoke subthreshold membrane potential responses. Labels indicate voltage responses recorded from the dendrite (Vd, grey) to somatic current (Is), and somatic responses (Vs, black) to dendritic current (Id). **B**) Scatter plot of normalized somato-dendritic transfer resistance (K_sd_) versus dendro-somatic transfer resistance (K_ds_) show that the transfer resistance is transitive. Transfer resistances were normalized to somatic input resistance (R_in_). Inset shows the overlay of traces from soma (black) and dendrite (grey) for the same family of current injections into the respective opposite compartment (accentuated in A). Note that they nicely match indicating transitivity. **C**) Relationship of normalized transfer resistance to distance of dendritic recording location from soma. The mono-exponential fit has a length constant of 383 μm. **D**) Scatter plot of K_ds_ normalized to K_ds_ measured at resting membrane potential versus the steady state depolarization at the dendritic recording site. K_ds_ values were estimated from the steady state depolarization in the soma to dendritic current steps of increasing amplitude in 38 neurons. Solid line indicates exponential fit (y=*Y0*\*e(*k*\*x)) to the data. **E**) Scatter plot of dendritic input resistance (K_dd_) normalized to K_dd_ measured at the resting membrane potential versus the local steady state depolarization. **F**) A simple two-compartment model was fitted to the recording data from individual neurons. Inset shows a schematic with a dendritic compartment (green) and a somatic compartment (red) connected by a resistor. The model included I_H_ (blue channels) and persistent sodium (yellow channel) and captured the nonlinear voltage-dependence of the recording data. For comparison, voltage responses to dendritic current steps in a linear 2-compartmental model without voltage-dependent conductances are shown in the top right corner. **G**) Scatter plot of K_ds_ normalized to somatic R_in_ (light grey) versus *D* determined from fits of the F-I data shows a systematic deviation from the identity line (*P*<0.0001, F(1,17)=94.74). If K_ds_ was measured during large current steps just below AP threshold (@depol; green), the slope is not significantly different from the identity line (*P*=0.27, F(1,17)=1.31), indicating a close match. Lines were constrained to cross the origin.

What are the mechanisms underlying the voltage-dependence of K_ds_? A simple linear two-compartment model without any voltage-dependent conductances cannot reproduce the observed behavior (Fig. 3F). A good candidate for mediating the underlying nonlinear conductance are HCN channels, which are strongly expressed in dendrites of pyramidal neurons (Magee, 1998; Williams & Stuart, 2000; Berger *et al.*, 2001; Harnett *et al.*, 2015). The HCN-mediated conductance is highly voltage-dependent with about half of the channels activated at −80 to −95 mV and an *e*-fold current response per 7 to 10 mV (Solomon & Nerbonne, 1993; Berger *et al.*, 2001). In neocortical L5 and CA1 pyramidal, HCN channels significantly contribute to the resting conductance in dendrites (Magee, 1998; Williams & Stuart, 2000; Berger *et al.*, 2001). As the reversal potential of the HCN-mediated current, I_H_, is close to −40 mV, I_H_ is the main driver for a more depolarized membrane potential in dendrites (Berger *et al.*, 2001). Including I_H_ in the two-compartmental model improved fits to the data (Fig. 3F). As expected, the fits showed that there was a high density of I_H_ specifically in the dendrites (135.4 ± 26.1 nS, versus 52.5 ± 8.8 nS in the soma, P=0.003, n=11 neurons). These results clearly demonstrated that voltage-dependent conductances in the somato-dendritic compartments of L5 pyramidal neurons are the basis for the strong voltage-dependence of the transfer resistance.

The voltage-dependence of the transfer resistance could potentially explain the large difference between the parameter *D* derived from the fits of the F-I relationship and K_ds_ normalized by R_in_ (Fig. 1L). K_ds_ had been measured on small dendritic current steps to minimize the influence of voltage-dependent nonlinearities. If K_ds_ was measured during large dendritic current injections inducing somatic depolarizations close to the AP threshold, K_ds_/R_in_ provided a much better match of *D* derived from the actual F-I relationship (Fig. 3G). Taken together, these results highlighted the synergistic interaction between depolarization in the somatic and dendritic compartments even at subthreshold membrane potentials independent of Ca^2+^ channel activation.

### GABA_B_R activated K^+^ channels diminish voltage-dependent increase of input and transfer resistance

GABA_B_R activation is well known to increase conductance through GIRK channels (Newberry & Nicoll, 1984; Palmer *et al.*, 2012) and has been reported to recruit additionally other K^+^ channels such as two-pore domain K^+^ channels in L5 pyramidal neurons and entorhinal stellate cells (Deng *et al.*, 2009; Breton & Stuart, 2017). To test whether an increase of the K^+^-mediated membrane conductance in the dendritic compartment was sufficient to cause the experimentally observed changes in dendritic input resistance and transfer resistance after the baclofen puff, we fitted the two-compartmental model to the membrane potential responses in the control condition for all experiments in Fig. 2C (Fig. 4A-B). We then refitted the models to the membrane potential responses during baclofen ejection with the dendritic membrane resistance and the resistance between both compartments as the only free parameters. While the inter-compartmental resistance remained unchanged (27.7 ± 5.7 versus 27.6 ± 5.1 MΩ, n=5 neurons), the fits resulted in a consistent increase in the local K^+^-mediated membrane conductance across all experiments (16.7 ± 3.6 versus 11.5 ± 2.2 nS) and were sufficient to induce the experimentally observed changes in the dendritic input resistance and transfer resistance (Fig. 4C-D). These results suggested that K^+^ channel activation in the dendrite was a major contributor to reduced voltage-dependence of input and transfer resistances during dendritic GABA_B_R activation.

**Figure 4.**
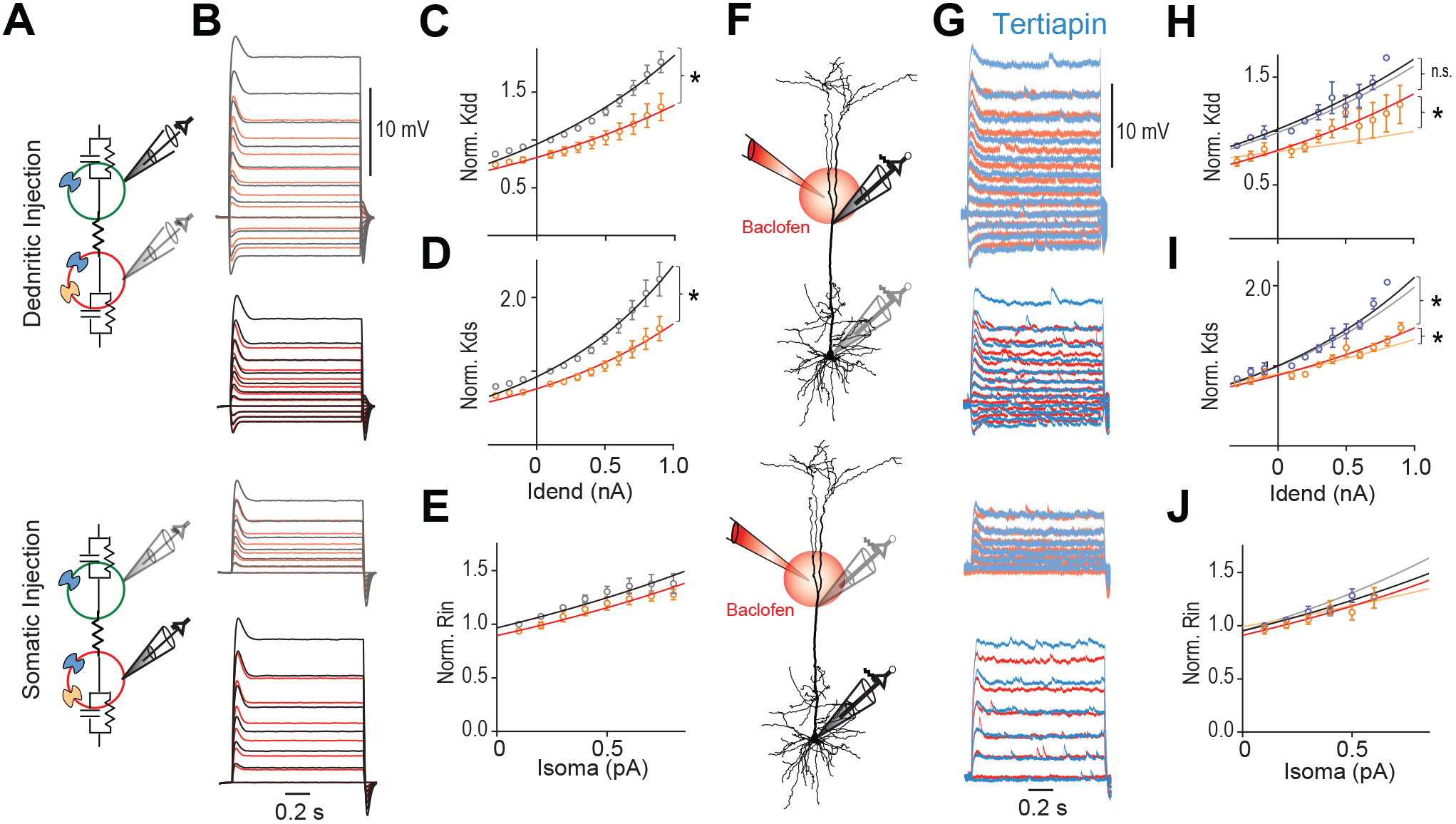
GABA_B_R-activated K^+^ channels reduce nonlinearities of input and transfer resistance. **A**) Two-compartmental models were fitted to the data of all experiments from Fig. 2C (n=5 neurons). **B**) Fitted voltage responses to families of current steps injected into either the dendritic (top) or somatic compartment (bottom) before (grey) and during the local application of baclofen (red) onto the apical dendrite. **C**-**E**) Increased K^+^-mediated conductance in the dendritic compartment was sufficient to capture the experimentally observed significant reductions of parameter *k* in the exponential fit (y=*Y0*\*e(*k*\*x)) of the voltage-dependence of K_dd_ (*P*=0.0166, n=5 models, F(1,116)=5.906) and K_ds_ (*P*=0.0348, F(1,116)=4.563) during the baclofen puff. **F**) Schematic of recording arrangement during dual dendritic and somatic patch-clamp recordings testing the contribution of GIRK channels. **G**) Families of current steps were injected into the dendrite (top) or soma (bottom) to evoke subthreshold membrane potential responses in the presence of tertiapin (0.5 μM, blue) and during the local application of baclofen (red) onto the apical dendrite. **H**-**J**) Tertiapin bath-applied partially blocked GABA_B_-mediated effects. Lighter shaded lines indicate the fitted curves in the absence of tertiapin from Fig. 2. The voltage-dependence of K_dd_ and K_ds_ was less affected by baclofen in the presence of tertiapin, as indicated by significantly larger *k*’s of the fitted lines (K_dd_: *P*=0.033, n1=4, n2=5, F(1,96)=4.678; K_ds_: *P*=0.0354, n1=8, n2=11, F(1,208)=4.484). In addition, the doubling interval of K_dd_ was no different after baclofen in the presence of tertiapin compared to tertiapin alone (2.2 nA vs. 1.9 nA; *P*=0.48, n=4 neurons, F(1,71)=0.5063).

### GIRK channel-activation in dendrites effectively inhibits action potential output

We tested experimentally the contribution of GIRK channels by applying the GIRK antagonist tertiapin (0.5 μM). Baclofen puffed onto the apical dendrite induced a small membrane potential hyperpolarization of −1.6 ± 0.2 mV and −0.7 ± 0.2 mV (n=15) in the apical dendrite and soma, respectively. Tertiapin reduced the baclofen-induced hyperpolarization by 54.1 ± 7.6% in the dendrite, consistent with a functionally significant contribution of GIRK channels to the baclofen-induced hyperpolarization (Extended Data Fig.4A). The incomplete block of the baclofen-induced hyperpolarization by tertiapin is in agreement with recent reports suggesting that other K^+^ channels such as two-pore domain K^+^ channels also contribute to GABA_B_-mediated membrane potential hyperpolarizations in L5 pyramidal neurons and entorhinal stellate cells (Deng *et al.*, 2009; Breton & Stuart, 2017). Importantly, tertiapin partially blocked the effect of baclofen on the voltage-dependence of input and transfer resistances (Fig.4F-J), suggesting that GIRK channel activation in the dendrite was an important factor that mediated the effect of dendritic GABA_B_R activation. We proceeded to test the effect of GIRK channel activation on the F-I relationship.

Bath application of tertiapin increased the somatic R_in_ (Extended Data Fig.4B). This resulted in greater cellular excitability reflected by increased values of the F-I gain (Extended Data Fig.4C). These observations suggested that there was a baseline activity of GIRK channels in the control condition. In the presence of tertiapin, the F-I relationship showed a strong nonlinear deviation from the PDSR model similar to the control condition (Fig. 5C). We proceeded to test the effect of the baclofen puff in presence of tertiapin (Fig. 5D-F). In the presence of tertiapin, baclofen was less effective in reducing AP output as shown by ANOVA analyses of the baclofen-induced AP rate reduction (Fig. 5G). Thus, there was a main effect of tertiapin (P=0.0131, F(1,6)=12.14, n =7), as well as a statistically significant interaction between tertiapin and stimulation-intensity (P=0.0083, F(3,18)=5.34). Post-hoc tests revealed that tertiapin reduced the effect of baclofen for all somatic current step sizes except 1000 pA, where the effect of baclofen was small anyway (Fig. 5G). Together, these results clearly demonstrated that dendritic GABA_B_R-mediated GIRK channel activation significantly contributed to reduced somatic AP output. The underlying mechanism was the modulation of the nonlinear integrative properties of the L5 pyramidal neuron rather than a pronounced impact on the somatic input resistance or membrane potential.

**Figure 5.**
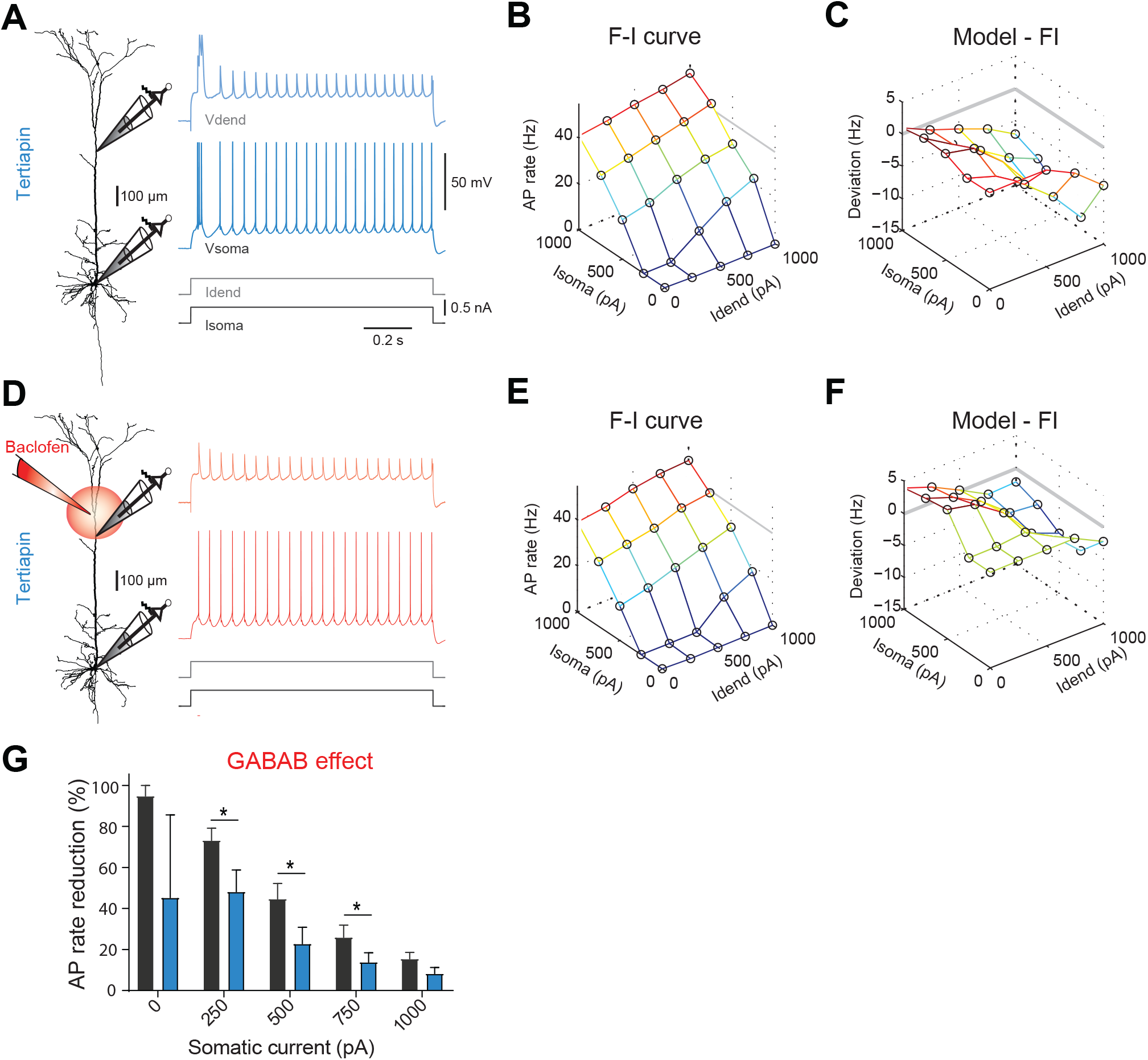
GIRK channel-activation in dendrites effectively inhibits AP output. **A**) Left, locations of dual dendritic and somatic patch-clamp recordings on a biocytin-filled layer 5 pyramidal neuron. Right, membrane potential responses of the same neuron to combined dendritic and somatic current injection during the presence of tertiapin (0.5 μM). **B**) Experimentally observed AP rate versus current (F-I) relationship for the same neuron. **C**) Average deviation of the experimental F-I relationship from the PDSR model across all neurons (n=7). **D**-**F**) Membrane potential responses and F-I relationship during a puff of baclofen in the presence of tertiapin. **G**) AP rate reduction during baclofen application in percent of baseline AP rate. Episodes of different dendritic current amplitude were grouped according to somatic current amplitude. There was a main effect of tertiapin (blue; P=0.0131, ANOVA, F(1,6)=12.14). Asterisks indicate significantly smaller reductions of the AP rate in the presence tertiapin (*P*<0.05, Bonferroni’s multiple comparisons test), demonstrating a block of the baclofen effect.

### The role of L-type Ca^2+^-channels in GABA_B_R-mediated inhibition

The previous results highlighted the importance of the nonlinear integrative properties of L5 pyramidal neurons at subthreshold membrane potentials. In how far was the F-I relationship shaped by dendritic Ca^2+^ spikes? To test this, we pharmacologically inhibited L-type Ca^2+^ channels by bath application of nimodipine (10 μM). Nimodipine inhibited dendritic Ca^2+^ spikes (cf. Fig. 6A vs. D) and hence reduced factor *D* of the F-I relationship (*P*=0.047, n=7, Wilcoxon signed rank test; Fig. 6L). However, the overall spike rate was not decreased (Fig. 6B vs. E) and the F-I relationship continued to significantly deviate from the PDSR model (Fig. 6C vs. F). Analysis of the inter-spike interval (ISI) distribution showed that nimodipine specifically reduced the number of fast ISIs of <15 ms emitted during Ca^2+^ spike-induced high frequency bursts (HFB) and resulted in an increased fraction of intermediate ISIs of 20 to 40 ms (Fig.6J). At the same time, the prominent slow afterhypolarzations (sAHP) following HFBs associated with particularly long ISIs (arrows in Fig.6A) also disappeared reflecting decreased activation of Ca^2+^ activated K^+^ channels in agreement with previous reports (Moyer *et al.*, 1992).

**Figure 6.**
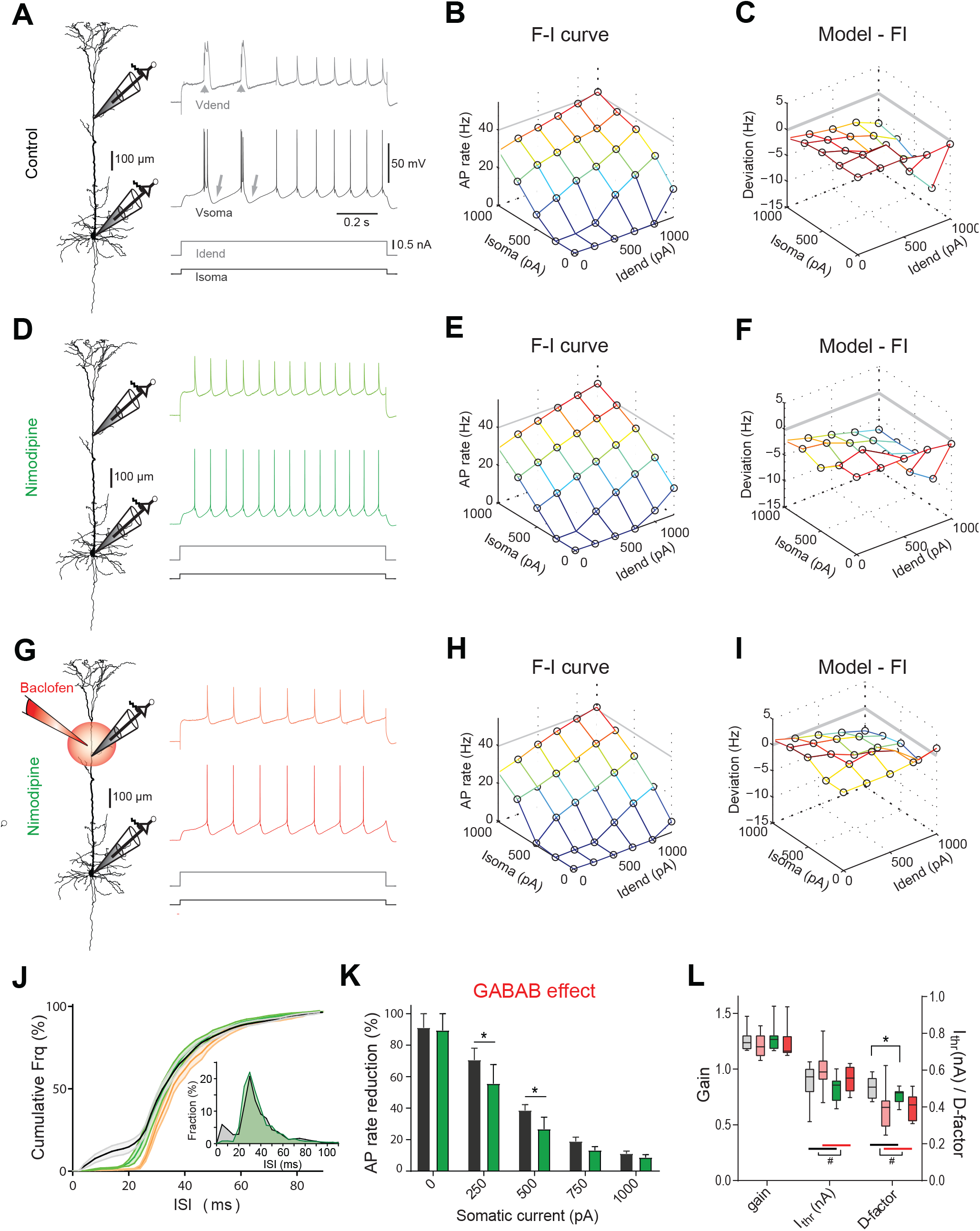
GABA_B_R-mediated inhibition of L-type Ca^2+^ channels prevents burst firing. **A**) Left, locations of dual dendritic and somatic patch-clamp recordings on a biocytin-filled layer 5 pyramidal neuron. Right, membrane potential responses of the same neuron to combined dendritic and somatic current injection. Dendritic Ca^2+^ spikes (arrowheads) and subsequent sAHPs (arrows) are indicated. **B**) Experimentally observed AP rate versus current (F-I) relationship for the same neuron. **C**) Average deviation of the experimental F-I relationship from the PDSR model across all neurons (n=7). **D**) Membrane potential responses of the same neuron to combined dendritic and somatic current injection after application of nimodipine (10 μM). **E**) Experimentally observed F-I relationship. **F**) Average deviation of the experimental F-I relationship during nimodipine application from the PDSR model across all neurons (n=7). Note the similarity to C. **G-I**) Membrane potential responses and F-I relationship during a puff of baclofen in the presence of nimodipine. Note the decreased deviation from the PDSR model. **J**) Grand mean of the cumulative ISI distribution (n=7 neurons). In control condition (grey), there was a high proportion of fast ISIs (<20 ms), which were lost after baclofen puff onto the apical dendrite (orange) or wash-in of nimodipine (green). Shaded area indicates the SEM. Inset shows the frequency distribution of all ISIs in control condition and after wash-in of nimodipine. Bin width was 5 ms. **K**) AP rate reduction during dendritic puff of baclofen in percent of baseline AP rate. Episodes of different dendritic current amplitude were grouped according to somatic current amplitude. Asterisks indicate significantly smaller reductions of the AP rate in the presence of nimodipine (green; *P*<0.01, Bonferroni’s multiple comparisons test), indicating a partial occlusion of the baclofen effect. **L**) Parameters of eq. (*2*) from fits to F-I data in control condition (gray) and in the presence of nimodipine (green) alone or in combination with a baclofen puff (light red and red; n=7). Main effects of baclofen (#, P<0.05, ANOVA) and significant differences between groups (*, P<0.01, Wilcoxon signed rank test) are indicated.

Next, we tested whether the effect of puffed baclofen was decreased in the presence of nimodipine (Fig. 6G-I). Similar to the control situation, baclofen reduced the overall AP output (Fig. 6H) and rendered the F-I relationship more similar to the PDSR model (Fig. 6I). Thus, puffed baclofen not only prevented HFBs but also shifted the whole ISI distribution towards longer ISIs (Fig.6J). ANOVA analyses of the spike rate reduction after the baclofen puff in the absence and presence of nimodipine within the same neurons did not show a main effect of nimodipine (P=0.12, F(1,6)=3.38), but revealed a significant interaction between nimodipine and stimulation intensity (P=0.0166, F(3,18)=4.45). Post-hoc tests revealed that the presence of nimodipine moderately decreased the impact of baclofen at intermediate somatic current levels (Fig. 6K).

Taken together, these results show that direct inhibition of L-type Ca^2+^-channels by dendritic GABA_B_Rs plays only a minor role in shaping the F-I relationship.

### Dendritic GABA_B_R activation decouples dendritic input from somatic AP output

The conclusions so far were derived from experiments involving linear current steps. Is the direct inhibition of L-type Ca^2+^-channels by dendritic GABA_B_Rs more prominent during more realistic time-varying input patterns? To test this, we injected current into the soma and dendrite in the shape of responses recorded *in vivo* during contralateral hind limb stimulation (Fig. 7A; Palmer et al., 2012). Combining these dendritic and somatic current waves at increasing amplitudes resulted in a complex pattern of evoked APs that included HFBs of two or more APs with an ISI of <15 ms. High-frequency bursts occurred preferentially during episodes with large dendritic current amplitudes at well-defined time points (Fig. 7B and C, Extended Data Fig. 7B). Under control conditions, HFBs were commonly associated with dendritic Ca^2+^ spikes, while APs separated by ISIs>15 ms rarely coincided with dendritic Ca^2+^ spikes (Fig. 7H). Activation of dendritic GABA_B_Rs by a puff of baclofen dramatically decreased HFB rate and eliminated Ca^2+^ spikes (Fig. 7D-E and I).

**Figure 7.**
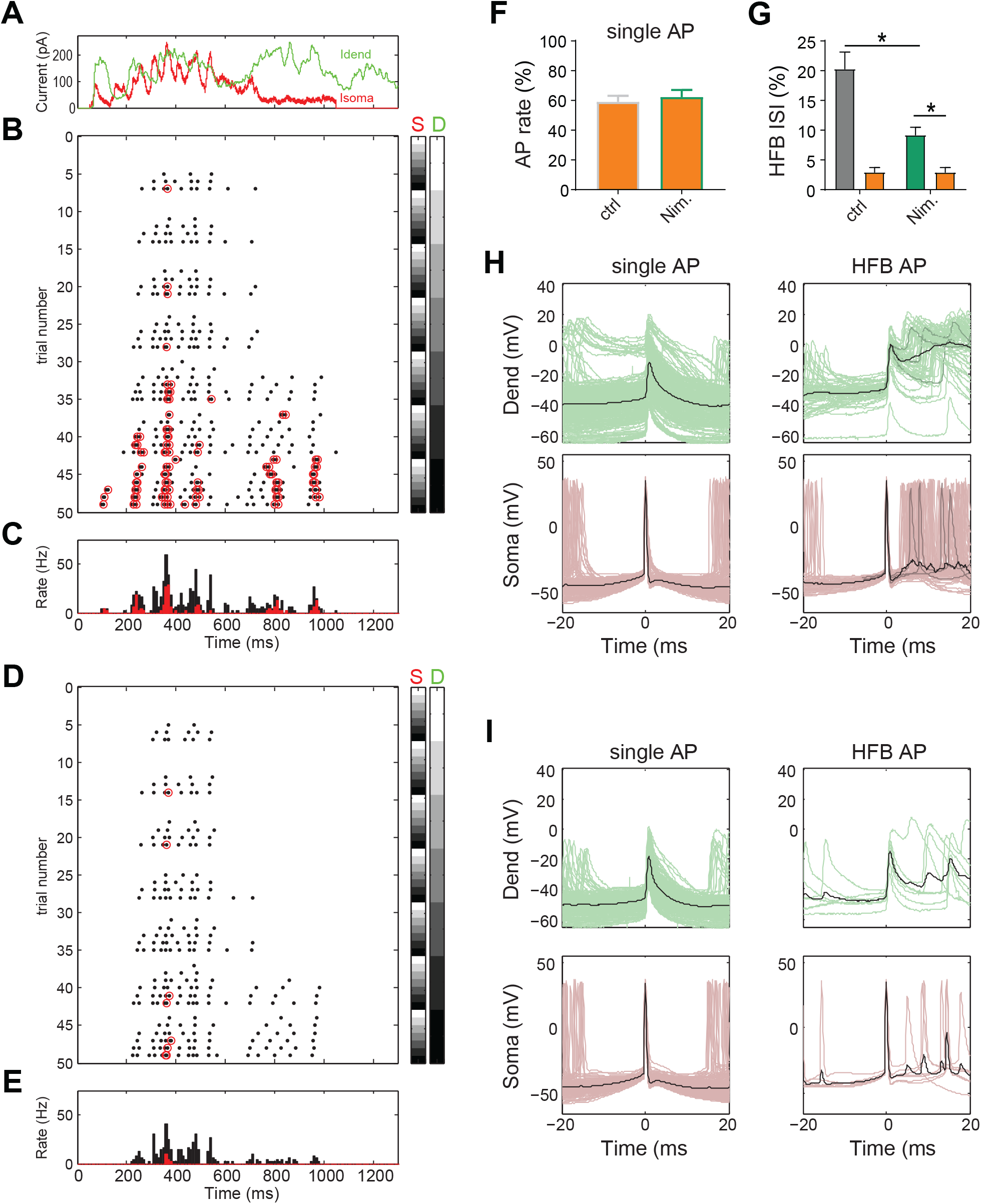
Differential regulation of single APs and HFBs during *in vivo*-like input waveforms by dendritic GABA_B_Rs. **A**) Injected current waveforms based on responses to sensory stimulation of the contralateral hind limb recorded *in vivo* from dendrite and soma, respectively (Palmer *et al.*, 2012). Dendritic current is shown in green, somatic in red. **B**) Raster plot of APs emitted in individual episodes during increasing levels of dendritic and somatic stimulation strength. The seven different levels of the injected current in 49 combinations are indicated by the right color bars (S=somatic, D=dendritic). The peak amplitude of the current waveform was increased from 0 pA (white) to 1500 pA (black). Step size was 250 pA. Red circles indicate APs that are part of a high-frequency burst (HFB, ISI<15ms). **C**) Peri-stimulus time histogram of all APs (black) and APs that are part of a HFB (red). **D**-**E**) Raster plot and PSTH of APs emitted in the same neuron while puffing baclofen onto the apical dendrite. **F**) Normalized AP rate during the dendritic baclofen puff in episodes of low AP probability (<7 Hz, and maximal dendritic current amplitude of 750 pA). The blockade of L-type Ca^2+^ channels by nimodipine had no effect on the effect of baclofen (P=0.64, n=7). **G**) The proportion of HFB ISIs (<15 ms) was significantly decreased by nimodipine (P=0.016; Wilcoxon Signed Rank test, n=7). Baclofen (orange) puffed onto the dendrite further decreased the proportion. **H**) Voltage traces recorded in dendrite (green) and soma (red) for all single APs outside of HFBs (ISI>15 ms, left) and APs at the start of a HFB (right). Three traces are highlighted to exemplify the variability of dendritic voltage traces during HFBs. **I**) Voltage traces recorded in dendrite and soma while puffing baclofen onto the apical dendrite.

What determined whether HFBs rather than individual APs were emitted in the two conditions? We analyzed the spike-triggered average (STA) of the dendritic and somatic current. To obtain a more accurate estimate of the current that effectively drove the axonal AP output, the dendritic current was scaled by *D* (I_eff_). *D* was obtained from the fit of *eq. 2* to the F-I relationship during linear current step injections in the same neurons under the same pharmacological conditions (Fig.1). The somatic I_eff_ was set equal to the injected somatic current. This analysis showed that, on average, a combination of constant intermediate dendritic I_eff_ of 271.3 ± 20.6 pA (n=7) and a peak somatic I_eff_ of 564.4 ± 42.4 pA drove single APs outside of bursts (ISI>15 ms; Extended data Fig. 7B). By contrast, HFBs were evoked by significantly higher levels of dendritic I_eff_ (P=0.016) in combination with somatic I_eff_ that was in the rising phase at the time of the first AP of the burst. The contribution of somatic (462.3 ± 33.2 pA, n=7) and dendritic I_eff_ (452.2 ± 35.6 pA) at the time of the first AP of the HFB was approximately equal, indicating that HFBs were often the result of the synergistic interaction between membrane depolarizations in the dendritic and somatic compartments in agreement with their close association to dendritic Ca^2+^ spikes.

The activation of dendritic GABA_B_Rs by baclofen altered the balance towards large somatic versus relatively small dendritic I_eff_ during initiation of single APs (somatic: 685.3 ± 48.2 pA vs. dendritic: 227.1 ± 27.5 pA). This effect was even more pronounced for APs at the start of the few remaining HFBs (somatic: 845.3 ± 33.4 pA vs. dendritic: 292.1 ± 23.3 pA), completely shifting the balance towards somatic I_eff_. This indicated that APs were only evoked at times of large somatic current, as stronger somatic current was required to compensate for less effective dendritic current.

Next, we directly tested the contribution of dendritic Ca^2+^ spikes to the initiation of HFBs. Pharmacological blockade of L-type Ca^2+^ channels by bath application of nimodipine (10 μM) decreased the relative proportion of HFB ISI from 20.3 ± 2.8% to 9.2 ± 1.3% of all ISIs (P=0.016; Wilcoxon Signed Rank test, n=7 neurons; Fig.7G). The contribution of dendritic I_eff_ to the remaining HFBs was significantly smaller than in control condition (359.9 ± 31.2 pA vs. 452.2 ± 35.6 pA; P=0.016), while significantly greater somatic current was necessary to evoke HFBs (600.1 ± 40.0 pA vs. 462.3 ± 33.2 pA; P=0.016; Extended data Fig. 7C). Direct activation of dendritic GABA_B_Rs by baclofen puffed onto the apical dendrite further reduced the HFB ISI proportion to 2.9 ± 0.8% comparable to the baclofen-induced effect in the absence of nimodipine (2.9 ± 0.8%). Thus, inhibition of L-type Ca^2+^ channels had qualitatively the same effect on HFBs as dendritic GABA_B_R activation.

By contrast, the inhibition of L-type Ca^2+^ channels did not change the contribution of dendritic I_eff_ to the induction of single APs outside of HFBs (P=0.81; Extended data Fig. 7B and C). This suggested that the reduction of the AP frequency in the absence HFB by dendritic GABA_B_Rs was independent of L-type Ca^2+^ channel inhibition. To directly test this hypothesis, we restricted the analysis to episodes with low AP frequency (<7 Hz). For this analysis, only episodes with an intermediate dendritic current level (max. 750 pA) were included to avoid any HFBs due to dendritic Ca^2+^ spikes. In episodes with low AP frequency, activation of dendritic GABA_B_Rs by puffed baclofen had the same effect in absence or presence of nimodipine (P=0.64, n=7; Fig.7F). This observation strongly supported the conclusion from experiments involving current step injections (Fig.4 and 5) that dendritic K^+^ channels are the main effectors of dendritic GABA_B_Rs to decrease the AP output.

Taken together, the results from both linear current steps and time-varying inputs converge at the conclusion that inhibition of L-type Ca^2+^ channels and activation of K^+^ channels by dendritic GABA_B_Rs synergistically alter the nonlinear integrative properties of L5 pyramidal neurons to suppress burst firing and decrease AP output under broadly varying conditions.

## DISCUSSION

By systematically mapping the input-output relationship during dual patch-clamp recordings from soma and apical dendrite, we demonstrated here for the first time that voltage-dependent local input and transfer resistances mediate a synergistic interaction between dendritic and somatic membrane depolarization. This was the principal mechanism causing a significant deviation of the F-I relationship from the predictions based on a model with a passive dendrite (PDSR). Dendritic GABA_B_R-activated K^+^ channels greatly reduced the voltage-dependent transfer resistance, while having negligible effects on the somatic input resistance and membrane depolarization at membrane potentials well below action potential threshold. This explains the surprising impact of interhemispheric inhibition on AP output in the absence of a discernable membrane hyperpolarization at the soma of L5 pyramidal neurons observed *in vivo* (Palmer et al., 2012). In contrast, the GABA_B_R-mediated direct block of L-type calcium channels had a more restricted effect and primarily prevented HFBs. These results highlight the powerful modulation of the input integration in pyramidal neurons by GABA_B_R-activated K^+^ channels.

### The significance of the voltage dependence of K_ds_

The contribution of dendritic K^+^ channels to the GABA_B_R-mediated response was much more pronounced than previously thought (Breton & Stuart, 2012; Palmer *et al.*, 2012). The main reason for the powerful impact of the seemingly small hyperpolarization by GABA_B_R-activated K^+^ channels (Newberry & Nicoll, 1984; Palmer *et al.*, 2012; Breton & Stuart, 2017) was due to its impact on other voltage-dependent conductances in the dendrites. While the local somatic and dendritic input resistances were weakly voltage dependent, the transfer resistance from dendrite to soma (K_ds_) strongly depended on the dendritic membrane potential doubling at an estimated rate of every 28.7 mV depolarization step (Fig.3D). Consequently, the ratio of K_ds_ to R_in_ increased and the impact of dendritic inputs on the somatic membrane potential grew supralinearly with increasing dendritic membrane potential depolarization. Incorporating K_ds_ measured at depolarized membrane potential provided a much more accurate estimate of the dendritic impact (*D*) on the somatic AP output.

The voltage-dependent deactivation of I_H_ appeared to be an important factor mediating the nonlinear K_ds_. At rest, I_H_ decreases the impact of small EPSPs evoked in the distal dendrite at the soma (Golding *et al.*, 2005; George *et al.*, 2009) and contributes to increased compartmentalization of synaptic inputs (Harnett *et al.*, 2015) that has also been observed in human L5 neurons (Beaulieu-Laroche *et al.*, 2018). However, larger depolarization deactivates I_H_ thereby increasing the local input resistance and decreasing the leakage for dendritic current flowing to the somatic compartment. This mechanism dramatically increases the impact of apical dendritic inputs on the soma when the dendrite is sufficiently depolarized.

### Mechanisms of GABA_B_R-mediated modulation of dendritic integration

Dendritic GABA_B_R activation greatly lowers the dendritic membrane resistance. Although the membrane hyperpolarization induced by the dendritic puff of baclofen appeared to be small (Breton & Stuart, 2012; Palmer *et al.*, 2012), the strong impact on the local input resistance (Fig.2C) suggests that the conductance change was much larger. GIRK channel activation locally hyperpolarizes the membrane potential and thereby recruits additional shunt by the activation of I_H_. In fact, the activation of I_H_ may have partially masked the actual hyperpolarization induced by GIRK channels. Interestingly, HCN channels appear to be structurally and functionally associated to GABA_B_Rs (Schwenk *et al.*, 2016). The combined effect of GIRK and HCN channel activation decreased the transfer resistance from dendrite to soma. Under these conditions, L5 pyramidal neurons behaved similar to the predictions of the simple PDSR model. In combination with the direct block of L-type Ca^2+^ channels (Perez-Garci *et al.*, 2013), GABA_B_R-activated K^+^ channels strongly down-modulate the impact of dendritic input currents on somatic AP discharge under all conditions.

### The presynaptic interneuron population responsible for dendritic GABA_B_R activation

GABA_B_R activation mediates a slow acting form of interhemispheric inhibition (Palmer *et al.*, 2012). Activation of the contralateral sensory cortex by hind limb stimulation *in vivo* causes a GABA_B_R-mediated inhibitory effect on spike activity in the ipsilateral cortex of up to 400 ms (Palmer *et al.*, 2012). Interhemispheric callosal fibers preferentially activate interneurons in the upper cortical layers including L1 (Palmer *et al.*, 2012). GABA_B_Rs are thought to be activated by spill-over of GABA from synaptic release sites (Scanziani, 2000; Kohl & Paulsen, 2010). In principle, any dendrite targeting interneuron population could activate dendritic GABA_B_Rs in L5 pyramidal neurons given that they release sufficient amounts of GABA to overcome the effective perisynaptic GABA reuptake mechanisms (Destexhe & Sejnowski, 1995; Thomson & Destexhe, 1999). However, cortical neurogliaform cells (NGFCs) appear to be of particular relevance for the modulation of dendritic properties via GABA_B_Rs. Thus, they are the only known interneurons that reliably activate postsynaptic GABA_B_Rs after a single presynaptic AP (Tamas *et al.*, 2003; Price *et al.*, 2008). Two anatomical features may contribute to the efficient GABA_B_Rs activation: the exceptional high density of about 1 bouton per 2.5 μm axon and the greater than usual distance of boutons from their target dendrites (Olah *et al.*, 2009; Overstreet-Wadiche & McBain, 2015). Hence, GABA released by NGFCs is thought to act via volume transmission potentially affecting many postsynaptic targets simultaneously rather than by ‘point-to-point’ synaptic transmission. Interestingly, NGFCs also engage a particularly slow form of ionotropic inhibition that reflects the slow GABA dynamics within the synaptic cleft as well as the slow receptor kinetics of high-affinity GABA_A_R in the postsynaptic membrane (Tamas *et al.*, 2003; Karayannis *et al.*, 2010; Schulz *et al.*, 2018). A recent study demonstrated that a large fraction of L1 interneurons that express Neuron-Derived Neurotrophic Factor (NDNF) inhibits postsynaptic pyramidal neurons by a combination of slow GABA_A_R and GABA_B_R-mediated IPSCs as it is typical for NGFCs (Abs *et al.*, 2018). Furthermore, NDNF+ interneurons effectively inhibited Ca^2+^ spikes evoked in the apical dendrite of L5 pyramidal neurons (Abs *et al.*, 2018). Therefore, NDNF+ interneurons and other NGFCs in the superficial cortical layers are the prime candidates to activate dendritic GABA_B_Rs in L5 pyramidal neurons.

### Converging modulatory pathways

NGFCs receive inputs from many afferent cortical and subcortical areas that project to L1 of the cortex (Abs *et al.*, 2018). In general, these inputs signal top-down information from higher order brain areas to control the processing of bottom-up information that is locally processed in the specific cortical area (Larkum, 2013). In addition, NGFCs also receive prominent neuromodulatory inputs such as cholinergic inputs from the basal forebrain (Lee *et al.*, 2010; Abs *et al.*, 2018; Poorthuis *et al.*, 2018). These observations show that modulation by NGFCs can be recruited via multiple converging associational pathways (D’Souza & Burkhalter, 2017; Schuman *et al.*, 2019).

Besides GABA, other neuromodulators including adenosine, serotonin, acetylcholine, endocannabinoids and somatostatin can activate GIRK channels in pyramidal neurons (Takigawa & Alzheimer, 1999; Seeger & Alzheimer, 2001; Chen & Johnston, 2005; Kim & Johnston, 2015; Lucas & Armstrong, 2015; Stumpf *et al.*, 2018). The results of the present study suggests that these neuromodulatory pathways will strongly affect dendritic integration.

The functional significance of dendritic GIRK channels has recently been highlighted by the observation that the higher induction threshold for LTP in dorsal versus ventral CA1 pyramidal neurons may be caused by tonically enhanced GIRK activation in dorsal CA1 pyramidal neurons (Malik & Johnston, 2017). In agreement with our results, increased GIRK channel activation in dorsal CA1 pyramidal neurons effectively reduces temporal integration of synaptic inputs associated with a diminished propensity to develop dendritic plateau potentials during bursts of pre- and postsynaptic activity (Malik & Johnston, 2017). In neurological disorders involving increased GIRK channel expression such as trisomy of chromosome 21 (Kleschevnikov *et al.*, 2012), increased dendritic GIRK channel activation is therefore expected to contribute to both deficits in dendritic integration and synaptic plasticity.

In conclusion, the results of the present study demonstrate that GABA_B_R-mediated activation of dendritic K^+^ channels effectively modifies the nonlinear integrative properties of L5 pyramidal neurons to decrease AP output under broadly varying conditions.

## METHODS

### In vitro preparation

Wistar rats (P30-P46) were anaesthetized with 95% CO_2_ / 5% O_2_ before decapitation. The brain was then rapidly transferred to ice-cold, oxygenated artificial cerebrospinal fluid (ACSF) containing (in mM) 125 NaCl, 25 NaHCO_3_, 2.5 KCl, 1.25 NaH_2_PO_4_, 1 MgCL_2_, 25 glucose and 2 CaCl_2_ (pH 7.4). Parasagittal slices of the primary somatosensory cortex (300 μm thick) were cut with a vibrating microslicer (Leica) and incubated at 37 °C for 30 minutes and subsequently maintained at room temperature (~22 °C). Dual somatic and dendritic whole-cell patch recordings were made from visually identified neurons using infared Dodt gradient contrast or oblique illumination and a CCD camera (CoolSnap ES, Roper Scientific). During recordings, slices were bathed in ACSF maintained at 33-35 °C.

### Patch-clamp recordings

Whole-cell patch-clamp recordings were obtained from thick-tufted layer 5b (L5) pyramidal neurons using a patch pipette (resistance 6-9 MΩ) filled with an intracellular solution containing (in mM): 135 K gluconate, 4 KCl, 10 mM HEPES, 10 Na_2_ phosphocreatine, 4 Mg-ATP, 0.3 Na-GTP, 0.2% biocytin, adjusted to pH 7.3-7.4 with NaOH. Dual whole-cell voltage recordings were performed from the soma and dendrite (resistance 20-40 MΩ) using Axoclamp 2A (Molecular Devices) and Dagan BVC-700A amplifiers (Dagan Corporation). Voltage was filtered at 5 kHz and digitized at 10 kHz using a BNC900 (National Instruments, Austin, TX) or ITC-18 interface (InstruTech, Port Washington, NY). Custom written igor software was used for acquisition. No correction was made for the junction potential between the bath and pipette solutions. Dual recordings were made from the soma and dendrite in current clamp mode. Current of varying amplitude was injected through either one or both electrodes simultaneously. Step currents were 1 s long, and in other experiments, two different voltage responses to contralateral-HS recorded from the soma and dendrite *in vivo* (Palmer *et al.*, 2012) were injected into the soma and dendrite, respectively.

### Drugs applications

Baclofen (50 μM) was pressure ejected from a glass pipette (tip diameter: 1 μm) placed 50–100 μm distal to the dendritic patch pipette (approx. 500-800 μm from the soma). The volume directly affected by the drugs pressure ejected from the puffing pipette was estimated to have a radius of approx. 100 μm, as measured by a test application of the fluorescent indicator Alexa 594 into a brain slice. Nimodipine (Sigma) was dissolved at 20 mM in DMSO on the day of the experiment. In experiments testing GIRK channel contribution, tertiapin (Sigma) was added to the bath ACSF (0.5 μM) and puff pipette (5.0 μM).

### Neuronal reconstruction

After recordings, neurons were prepared for biocytin reconstruction. The slices were placed in 4% paraformaldehyde after the experiment for up to 4 days. Slices were processed for biocytin staining to reveal the morphology of the recorded neuron. Neuronal reconstruction was then performed with Neurolucida software.

### Data analysis

Data were analyzed offline using MATLAB 7.13 with Signal Processing 6.16 and Statistics 7.6 Toolboxes. Action potentials (APs) were detected using a threshold criterion when the membrane potential crossed 0 mV. The time of the maximal depolarization was saved for each AP. The spike rate was defined as number of APs during the 1 s-long current step. Interspike intervals (ISI) were calculated as the difference in time between two subsequent APs. For experiments with fluctuating current stimuli, segments of the injected current waveform around the time of each AP were saved for visualization. APs were classified into three categories: single APs (preceding ISI_n-1_ and subsequent ISI_n_>15 ms), APs at the start of a burst (ISI_n-1_>15 ms and ISI_n_<15 ms), and APs during a burst (ISI_n-1_<15 ms). The spike-triggered average (STA) was calculated as the mean of these current segments over all spikes of a defined class. For the analysis of the baclofen-induced effect on trails with low spike rates (Fig.7F), only episodes with a maximal dendritic current amplitude of 750 pA and less than 7 spikes at baseline were included.

Input and transfer resistances were determined from the slope of a regression line fitted to four mean membrane potentials produced by a series of subthreshold current pulses around resting membrane potential (−100, 0, +100, +200 pA). For transfer resistances, current was injected into either somatic or dendritic electrode and membrane potentials were measured at the opposite electrode. For experiments involving tertiapin application, K_ds_ and R_in_ were directly derived from 250 pA current injections into dendrite and soma, respectively. To test for a voltage-dependence of the input (transfer) resistance, input (transfer) resistances were determined for increasing current step amplitudes according to Ohm’s law and then normalized by the input (transfer) resistances measured at rest. For this analysis, only experiments with optimal compensation of series resistances were included.

#### Spike rate model

The frequency-input (F-I)-curve of somatic current injections was well fitted by a square root function (Fig. 1B):

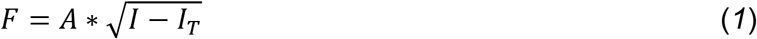

for all I>I_T_; where I_T_ denotes the current threshold and parameter *A* determines the slope, i.e. the gain of the input-output function. To predict the effect of additional current injection into the apical dendrite, we extended this spike rate model to include a scaled contribution of the dendritic current:

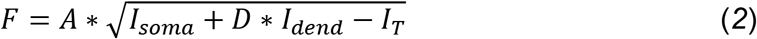

for all I>I_T_; where the factor *D* scales the relative impact of the dendritic current injection (I_dend_). Eq. (*1*) was fitted to AP rate data in response to increasing amplitudes of 1 s long current steps into the soma (Fig. 1B; 100 pA step size) using the Matlab function fminsearch. *A* and *I*_*T*_ were free parameters. Estimates of *A* and *I*_*T*_ were combined with an estimation of *D*=K_ds_/R_in_ to predict the AP rate of combined dendritic and somatic current injections according to the passive dendrite spike rate (PDSR) model. The parameters of the PDSR model were kept constant and were compared to the parameters derived from fits of recorded F-I relationship to eq. (*2*) during different pharmacological conditions. For fits of the AP rate data during combined somatic and dendritic current steps (Fig. 1E; 250 pA step size) to equation (*2*), *A*, *I*_*T*_ and *D* were free parameters. For experiments involving tertiapin application, parameter *A* of the PDSR model was directly derived from the fit to the AP rate data in response to combined dendritic and somatic current injections. The predicted spike rate from this model was compared to the measured spike rate in the presence and absence of an additional baclofen puff onto the apical dendrites.

#### Regression analyses

Values for *D* obtained from fits to the experimental F-I data were compared to theoretically predicted values using nonlinear regression analysis in GraphPad Prism 6. A regression line was fitted to the scatter plot of predicted versus measured values (Fig. 3G). The Y-axis intercept was constrained to zero. An extra sum-of-square F test was used to test for a significant deviation of the fitted slope from 1 indicating that theoretically predicted values systematically deviated from values derived from fits. Similarly, exponential functions were fitted to scatter plots of normalized input (transfer) resistances versus membrane potential or current step amplitude. Extra sum-of-square F tests were used to statistically compare growth rates between different data sets.

#### Two-compartment model

Two compartment models were simulated in the Python-based simulation environment Brian (Goodman & Brette, 2009). Membrane resistances of dendritic and somatic compartment as well as the resistance of the connecting resistor for linear models were derived from electrophysiological measurements of somatic R_in_ (K_ss_), dendritic R_in_ (K_dd_) and transfer resistance K_ds_ according to the following formulas. Dendritic membrane resistance:

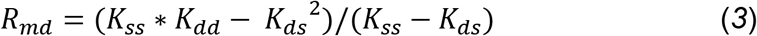

Axial resistance:

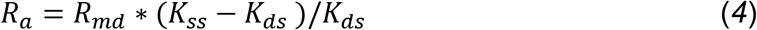

Somatic membrane resistance:

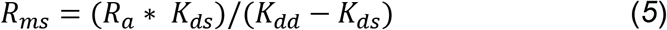

The capacitance for each compartment was estimated from the exponential fit to the decaying phase of small positive current steps at the soma and dendrite, respectively.

To model the voltage-dependence of the transfer resistance, persistent sodium channels were included in the somatic compartment, and HCN (hyperpolarization-activated cyclic nucleotide-gated cation) channels mediating I_H_ were included in both compartments. Persistent sodium current was modeled as:

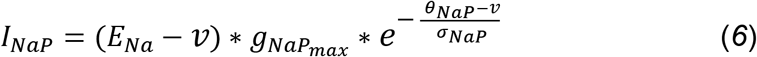

θ_NaP_ was set to −57.9 mV, and σ_NaP_ was set to 6.4 mV (Amarillo *et al.*, 2014).

I_H_ was modeled as:

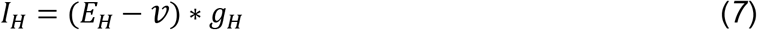

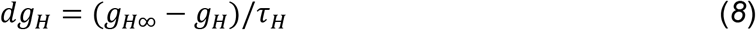

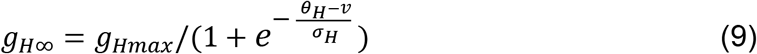

θ_H_ was set to −80 mV, σ_H_ to 12 mV, and τ_H_ to 40 ms, similar to published values in the literature (Solomon & Nerbonne, 1993; Berger *et al.*, 2001). The density of HCN and persistent sodium channels was derived from fits of experimentally observed steady-state membrane potential in soma and dendrites in response to somatic and dendritic current steps. The fitting procedure minimized the deviation of the model’s steady state voltage responses to the applied current steps from the experimentally observed steady-state membrane potential responses by adjusting six conductance densities: the somatic and the dendritic leak membrane conductance, the axial conductance (1/ R_a_), somatic and dendritic g_Hmax_ and somatic g_NaPmax_. Initial values for the first three parameters were the inverse of R_ms_, R_md_ and R_a_ calculated for the linear model (eqs. *3*-*5*). Initial values for dendritic and somatic g_Hmax_ and somatic g_NaPmax_ were set to 1/ R_md_, 1/ 20*R_ms_ and 1/ 5*R_ms_, respectively. Minimization was performed by the function minimize from the scipy.optimize package using the Sequential Least Squares Programming optimization algorithm (SLSQP). For fits of membrane potential responses during baclofen ejection, dendritic leak membrane conductance and axial conductance were the only free parameters, while the other four parameters were fixed at the values fitted to control responses.

#### Experimental Design and Statistical Analysis

Statistical analyses were performed in GraphPad Prism 6. All pharmacological tests were within experiment comparisons, i.e. the baclofen-induced effect in the presence of nimodipine/tertiapin were compared with the baclofen-induced effect alone in the same cell. The sample size was based on our experience with the high reproducibility of similar experiments (Palmer *et al.*, 2012). Data sets were analyzed with the nonparametric Wilcoxon Signed Rank and the Mann-Whitney tests for paired and unpaired data, respectively. The effect of nimodipine/tertiapin on the baclofen-induced AP frequency reduction was tested by a two-way ANOVA on normalized AP rate data. The significance level was set to P=0.05. All data are shown as mean ± s.e.m.. Unless stated otherwise, the number (n) of observations indicated reflects the number cells recorded from.

## Acknowledgements

We would like to thank Lucy Palmer and Enrique Perez-Graci for helpful comments on the manuscript and Natalie Nevian for technical assistance. This work was supported by the Swiss National Science Foundation (SNSF, PP00A‑102721 and 31003A_130694).

---

**Extended Data Figure 4.**
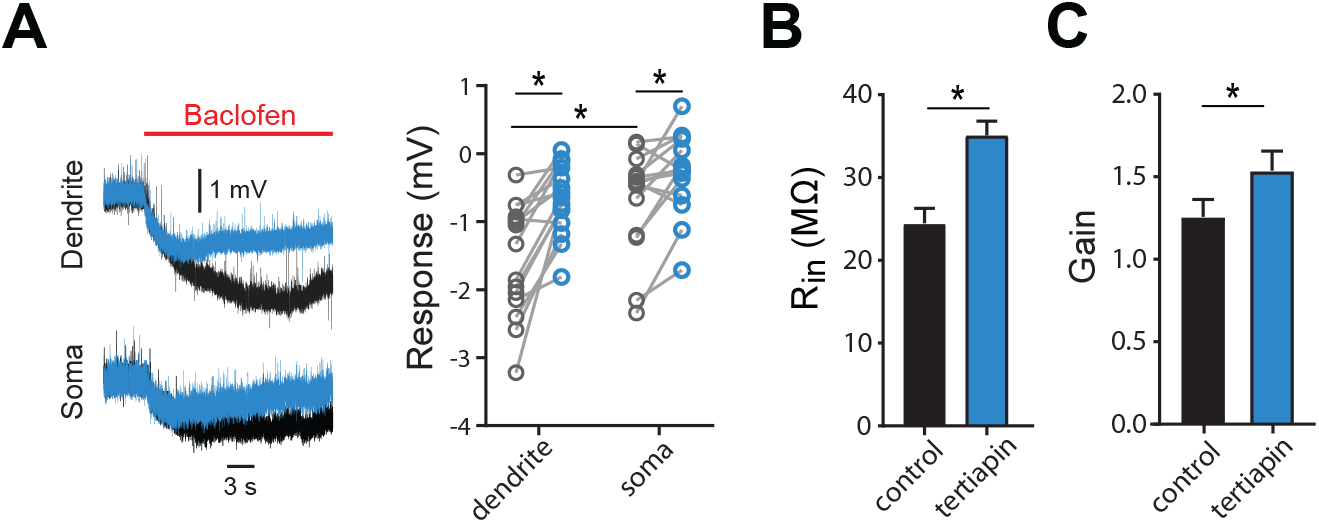
GABA_B_R-activated K^+^ channels reduce nonlinearities of input and transfer resistance. **A**) Local hyperpolarization in the dendrite and soma induced by the dendritic baclofen puff and partial block by tertiapin. Significant effects are indicated (*P*<0.05, n=15, Wilcoxon signed rank test). **B**) The somatic input resistance was significantly increased in the presence of tertiapin (*P*=0.0156, n=7, Wilcoxon signed rank test). **C**) The F-I gain derived from the fit of eq. (*2*) to the data was also increased (*P*=0.0156, n=7).

**Extended Data Figure 7.**
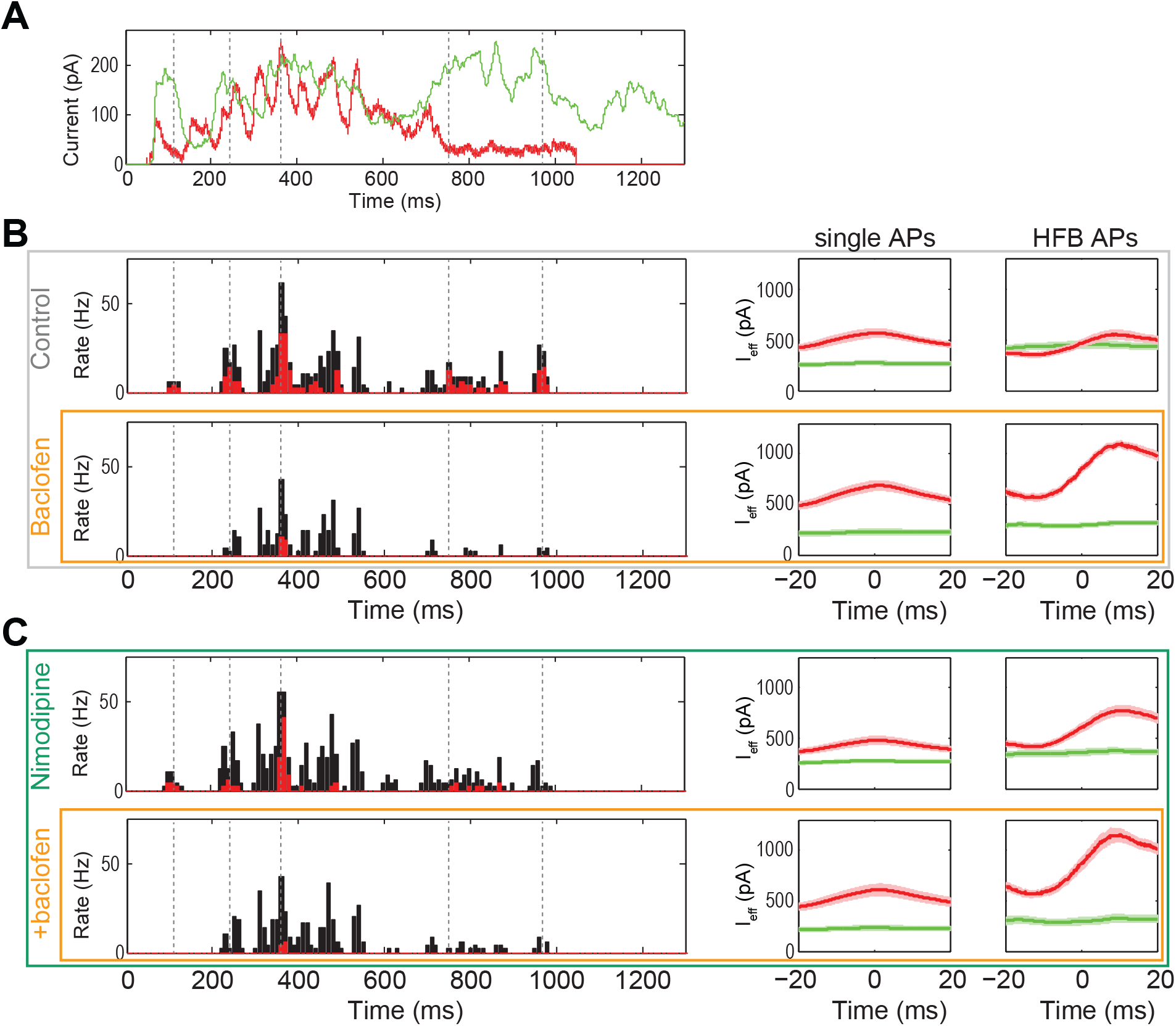
Differential regulation of single APs and HFBs during *in vivo*-like input waveforms by dendritic GABA_B_Rs. **A**) Injected current waveforms based on responses to sensory stimulation of the contralateral hind limb recorded *in vivo*. Dendritic current is shown in green, somatic in red. **B**) Peri-stimulus time histograms (PSTH, left) of all APs (black) and APs that are part of a HFB (red) in a neuron during control condition (top) and while puffing baclofen onto the apical dendrite (bottom). Vertical dashed lines in the PSTH and the current waveforms (A) indicate time points of consistently reoccurring HFBs across episodes. On the right, the grand mean (n=7 neurons) of the dendritic (green) and the somatic (red) I_eff_ is shown. Note the different temporal modulation of somatic I_eff_ and the increased level of dendritic I_eff_ necessary for the induction of HFB compared to single APs during the control condition. Dendritic activation of GABA_B_Rs by puffed baclofen strongly increased the contribution of somatic I_eff_, suggesting that AP output was mainly driven by the somatic current injection. **C**) The effect of the L-type Ca^2+^ channel blocker nimodipine on AP output. Left, PSTH from the same neuron before and during dendritic baclofen application in the presence of the L-type Ca^2+^ channel blocker nimodipine. Right, grand means of I_eff_ from the same group of neurons before and during baclofen application. Note the reduced contribution of dendritic I_eff_ and the increased contribution of somatic I_eff_ to HFBs in the presence of nimodipine.

